# Enhancing Retrieval Capacity of the Predictive Brain through Dorsolateral Prefrontal Cortex Intervention

**DOI:** 10.1101/2024.07.10.602829

**Authors:** Laura Szücs-Bencze, Teodóra Vékony, Orsolya Pesthy, Krisztián Kocsis, Zsigmond Tamás Kincses, Nikoletta Szabó, Dezso Nemeth

## Abstract

The ability to extract spatial or temporal regularities across experiences is crucial for skill development and predictive processes. The prefrontal cortex (PFC) plays a key role in modulating competitive memory systems, supporting declarative/episodic memory as opposed to statistical learning. This regulatory role may explain findings of improved acquisition and consolidation of statistical regularities following the suppression of dorsolateral PFC (DLPFC) by repetitive transcranial magnetic stimulation (rTMS). This raises a key question: Is access to models and prior statistical knowledge also modulated by the DLPFC? This preregistered study provides new insights by examining the role of the DLPFC in retrieving pre-existing knowledge of temporally distributed statistical regularities. Using a probabilistic learning task, healthy participants engaged in implicit statistical learning for 25 minutes. After a 24-hour consolidation period, participants received either 1 Hz rTMS or sham stimulation over the left, right, or bilateral DLPFC for 10 minutes before retesting. We found more effective access to statistical regularities in the bilateral DLPFC group compared to the sham group. Our results suggest that DLPFC suppression enhances the retrieval of statistical knowledge, particularly when interhemispheric compensatory mechanisms are prevented. These findings contribute to understanding competitive memory systems and offer implications for cognitive enhancement strategies.

## Introduction

In the dynamic interplay of multiple memory systems, the prefrontal cortex (PFC) emerges as a pivotal orchestrator, controlling the—sometimes competitive—interaction between distinct memory systems (Daw et al., 2011; Smalle et al., 2022; Smittenaar et al., 2013). Particularly, it is suggested to modulate the activity of the hippocampus and striatum, thereby influencing the switch between statistical learning and episodic memory, as well as habitual and goal-directed behavior (Oehrn et al., 2018; Sherman, Huang, et al., 2024; Sherman, Turk-Browne, et al., 2024; Sherman & Turk-Browne, 2020; Williams, 2023). In a neuroimaging study, the tract integrity between the hippocampus and dorsolateral prefrontal cortex (DLPFC) predicted the extent of statistical learning, which forms the basis for predictive processes and the construction of predictive models, providing further support for the role of the interplay between these two brain regions (Bennett et al., 2011). Nevertheless, PFC is known for its role in executive functions and cognitive control, processes that are proven to have an antagonistic relationship with habitual processes like statistical learning (Pedraza et al., 2024; Virag et al., 2015). Various methods of suppressing PFC activity, such as hypnosis (Nemeth et al., 2013), transcranial magnetic stimulation (TMS) (Ambrus et al., 2020; Smalle et al., 2022), or cognitive overload (Smalle et al., 2022), have been shown to—seemingly counterintuitively—enhance statistical learning performance. Neuroimaging connectivity studies also prove that the intricate neural background of statistical learning involves the disengagement of PFC circuitry (Park et al., 2022; Tóth et al., 2017), further indicating an antagonism between top-down control processes and statistical learning. This then raises the key question: Is access to models and prior acquired statistical knowledge also modulated by the DLPFC? While our understanding of the relationship between predictive model development and the PFC’s involvement is extensive, the mechanisms underlying the retrieval of previously learned statistical information remain largely elusive (Szücs-Bencze et al., 2023).

Non-invasive brain stimulation (NIBS) techniques, such as TMS, offer a means to explore causal relationships between brain regions and cognitive functions. The findings of the handful of studies that explored the impact of DLPFC modulation by TMS on statistical learning are somewhat contradictory (Szücs-Bencze et al., 2023). Some early studies reported diminished capacity for statistical learning following DLPFC neuromodulation (Pascual-Leone et al., 1996; Robertson et al., 2001), whereas others found better statistical learning due to DLPFC disruption with TMS (Ambrus et al., 2020; Galea et al., 2010). Additionally, the acquisition of linguistic regularities was also successfully enhanced by a patterned inhibitory TMS protocol over the DLPFC (Smalle et al., 2017, 2022). Nonetheless, certain studies failed to detect any influence of DLPFC stimulation on statistical learning (Gann et al., 2021; Wilkinson et al., 2010). In those studies where reduced statistical learning performance or no effect was observed following DLPFC stimulation, either high-frequency repetitive TMS (rTMS), likely inducing heightened excitability rather than inhibiting DLPFC (Pascual-Leone et al., 1996), or inhibitory TMS protocols were applied, but prior to the learning phase (Gann et al., 2021; Robertson et al., 2001; Wilkinson et al., 2010). Conversely, successful boosting of statistical learning was noted in studies utilizing inhibitory TMS protocol timed after the learning phase, thus promoting enhanced consolidation and subsequent knowledge retention (Ambrus et al., 2020; Galea et al., 2010; Tunovic et al., 2014). Beyond the scope of learning and acquisition, the application of probabilistic knowledge and predictive processes is a continuous necessity in daily life. For peak cognitive performance, the processes require consistent access and retrieval. This prompts the intriguing inquiry: might the modulation of the DLPFC impact the retrieval processes as profoundly as it does the acquisition ones?

In order to fill this gap, our study aimed to investigate the effect of inhibitory DLPFC stimulation on the retrieval of pre-existing knowledge of statistical regularities. Brodmann 9 was chosen as the target site as previous studies have shown boosting effects on statistical learning and predictive processes (Ambrus et al., 2020; Galea et al., 2010; Smalle et al., 2022). The comprehensive mapping of the role of DLPFC was taken by targeting the left, right, and bilateral DLPFC in separate groups with low-frequency rTMS. We applied bilateral stimulation, which is a unique and surprisingly rarely used method in cognitive neuroscience research, however, this design is assumed to minimize compensatory mechanisms by the non-stimulated hemisphere. Moreover, as recent studies indicate that it is unjustified to determine TMS intensity based on the motor threshold when stimulating areas outside the primary motor cortex (Antal et al., 2004; Turi et al., 2022; Wassermann et al., 1992), we employed a uniform intensity setting, similarly to Ambrus et al. (2020). The fixed intensity was determined by simulating the electric field in the brain, using SimNIBS 4 (Thielscher et al., 2015). We evaluated implicit statistical learning by the Alternating Serial Reaction Time (ASRT) task (Howard et al., 2004), which reflects real-world statistical learning processes with high reliability (Farkas et al., 2023). To control the specificity of rTMS on the retrieval of statistical knowledge, we used the Paired Associate Learning Task (PALT) as a control task, measuring declarative learning and recall. After participants learned on the tasks, a 24-hour retention period ensued. Subsequently, prior to retesting (Retrieval session), participants received inhibitory stimulation in the form of 1 Hz rTMS, or sham stimulation for 10 minutes. We have formulated hypotheses for all possible outcomes. If DLPFC modulation affects access to long-term memory representation and cognitive models, a decrease in both statistical and declarative retrieval is expected. However, if DLPFC stimulation weakens control functions, intact statistical retrieval alongside decreased declarative recall should emerge. Lastly, if DLPFC stimulation does not impact the retrieval of implicit statistical learning, it suggests that this form of learning is inherently automatic and robust, impermeable to disruption by rTMS following a lengthy consolidation period. Bilateral stimulation has already been proven to be effective in statistical learning modulation and achieving better learning outcomes (Ambrus et al., 2020). Based on this, we further anticipate that bilateral stimulation will yield the greatest benefit by preempting any potential interhemispheric compensatory mechanisms that may occur in the left and right DLPFC groups.

## Materials and methods

### Participants

One hundred and four healthy adult volunteers were enrolled in this preregistered study (https://osf.io/ns2km/). Two participants were excluded because it was revealed they had a neurological/psychiatric disorder, and one participant did not complete the study. Thus, the final sample consisted of 101 participants. They all had normal or corrected-to-normal vision and no TMS contraindications (pacemaker, history of major surgery, history of neurological or psychiatric disease, metal implant, pregnancy). None of the participants withdrew from the participation owing to TMS discomfort. The participants were randomly assigned to one of four groups: Left DLPFC, Right DLPFC, Bilateral DLPFC, or Sham group with 25, 26, 25, and 25 participants in each group, respectively (see Table 1 for descriptive statistics). Written informed consent was obtained from each subject. All study procedures were approved by the Regional Scientific and Research Ethics Committee of Albert Szent-Györgyi Clinical Center, University of Szeged, which complied with the Declaration of ‘Helsinki’s ethical guidelines.

**Table 1.**
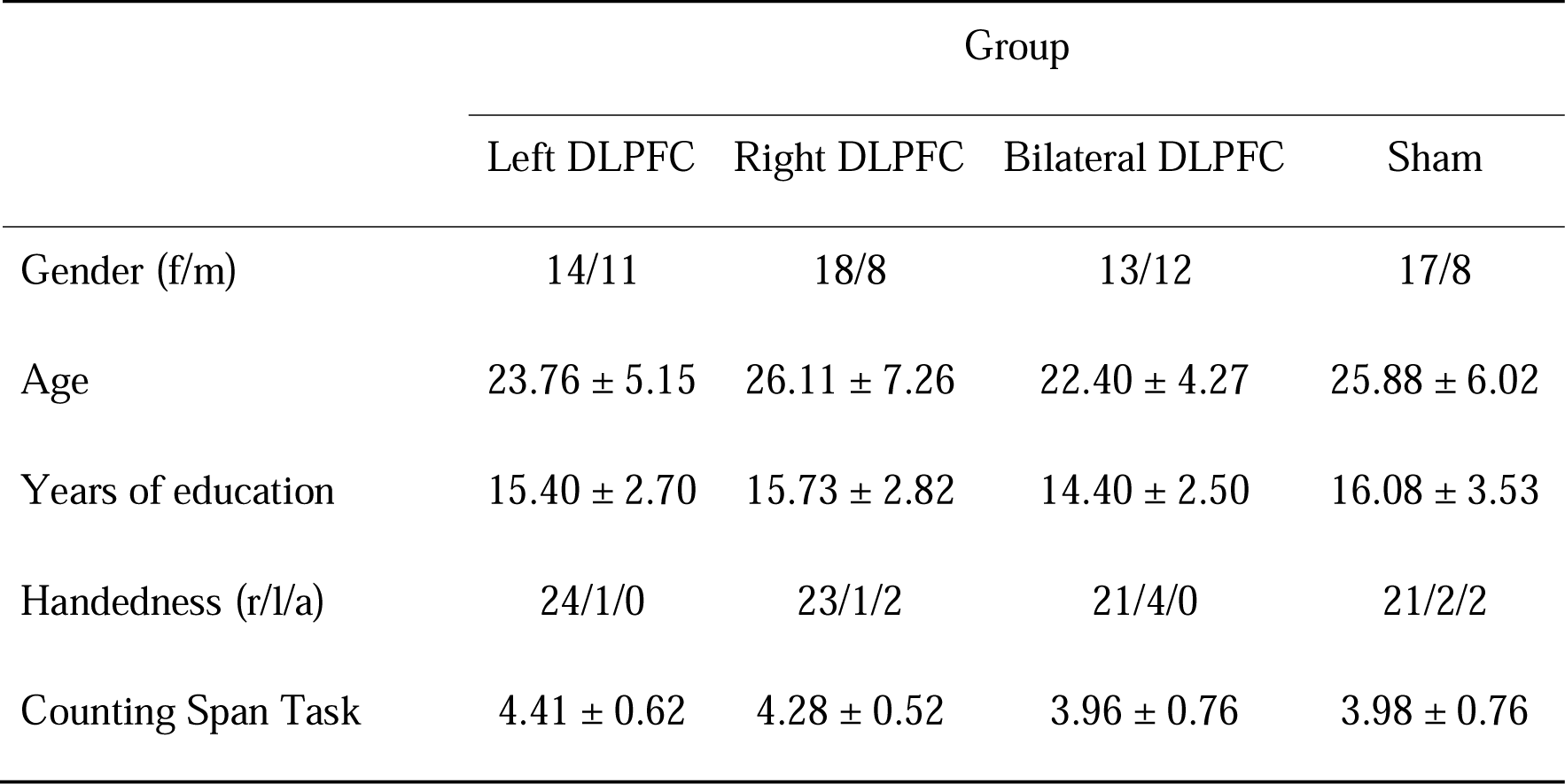
Descriptive statistics of the four experimental groups. *Note*. Mean and SD values for age, years of education, and Counting Span Task are presented. For gender (f = female, m = male), and handedness (r = right, l = left, a = ambidexter), case numbers are presented.

### ASRT task

Statistical learning was assessed via the ASRT task (Howard et al., 2004). Participants were presented with a stimulus (depicting a ‘dog’s head) appearing in one of four empty circles arranged horizontally on the screen (see Figure 1a). Their task was to press the corresponding keys (Z, C, B, or M on a QWERTY keyboard) swiftly and accurately. The participants were instructed to use their middle and index fingers of the left hand to press the Z and C keys, and the same fingers of the right hand to press the B and M keys, respectively. The stimulus remained on the screen until the subject pressed the correct key, after which the next stimulus appeared 120 ms later (response-to-stimulus interval). Unbeknownst to the participants, the stimuli followed a probabilistic eight-element sequence, where pattern and random elements alternated (e.g., 2r4r3r1r, where numbers 1-4 denoted target locations from left to right, and ‘r’ represented a randomly selected position of the four possible ones) (see Figure 1b). The task comprised 25 blocks, each with 80 trials, repeating the eight-element sequences ten times within each block. Due to the alternating pattern, the ASRT task provided the occurrence of certain sets of three consecutive stimuli (referred to as triplets) with varying probabilities. In high-probability triplets, the third element could be predicted based on the first element with higher probability (constituting 62.5% of all trials) compared to low-probability triplets, where the probability of the prediction of the third element from the first one was lower (constituting 37.5% of all trials) (see Figure 2b). We categorized each trial in a sliding window manner based on whether it represented the third element of a high-probability or low-probability triplet. Statistical learning was defined as the reaction time (RT) difference between trials that were the third element of a high-probability triplet or a low-probability triplet. Besides, participants generally become faster on the task irrespective of triplet types, indicating general visuomotor performance.

**Figure 1.**
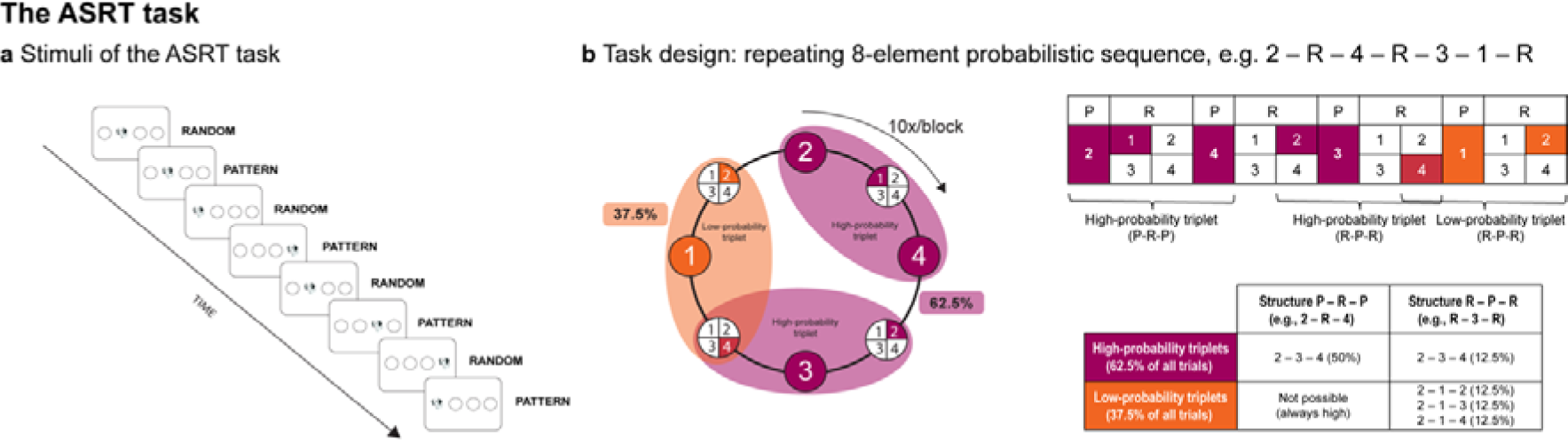
The ASRT task. **a** Pattern elements alternate with random elements. **b** An 8-element probabilistic sequence repeats ten times during a block. Due to this sequence structure, some runs of three successive stimuli appeared with higher probability (high-probability triplets) than others (low-probability triplets). Each trial was categorized as the last element of a high- or low-probability triplets. The RT difference between the two trial types indicates implicit statistical learning.

**Figure 2.**
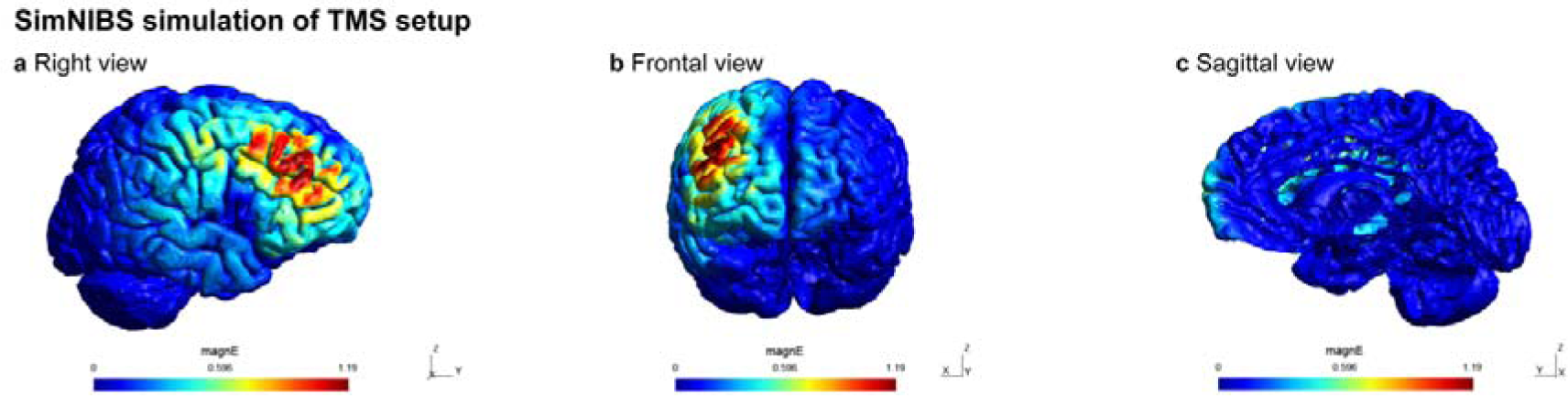
SimNIBS simulation of TMS setup. Demonstration of the spread of electrical field using SimNIBS 4 (“Earnie” head mash) when positioning the TMS coil over the right DLPFC (F4). Results are similar for the left DLPFC (F3) stimulation. SimNIBS did not allow us to simulate sequential bilateral stimulation. The electric field is represented as volt per meter (V/m).

### Control Memory Task

The PALT (Nagy et al., 2013) measuring declarative learning was utilized as a control memory task. During the learning phase of the PALT, participants were presented with pairs of images. Participants had to name the images they viewed and memorize them. During the recall phase of the PALT, participants encountered pairs of images again, some of which were presented with their original pairs (Old-Old original), while others were presented as rearranged pairs (Old-Old rearranged). Additionally, images that were not previously shown (New-Old or New-New) were included in the stimuli. Participants were required to determine whether they had seen these images before and, if so, whether they appeared together or separately during the learning phase.

### TMS Protocol

TMS stimulation was administered through a Magstim Rapid2 Stimulator equipped with a D702 70 mm figure-of-eight coil (The Magstim Company Ltd, Whitland, Wales, UK). Magnetic pulses were delivered at 1 Hz for 10 minutes, resulting in a total of 600 pulses. To determine our TMS setup, we used SimNIBS 4 (Thielscher et al., 2015). We demonstrate the results of the right DLPFC stimulation in Figure 2 as similar results were found in case of the left DLPFC stimulation. The stimulation intensity was uniformly set for all participants at 55% of the maximum stimulator output (MSO). We opted against the traditional motor threshold-based intensity setting as evidence indicated that this approach is inappropriate for stimulating regions outside the motor cortex (Antal et al., 2004; Turi et al., 2022; Wassermann et al., 1992). For instance, motor thresholds can vary significantly even among different upper extremity muscles (Wassermann et al., 1992). Moreover, there is no correlation between motor threshold and TMS-induced effects in other cortical areas, such as the induction of phosphenes in the visual cortex (Antal et al., 2004). These findings suggest that using motor threshold as a basis for determining stimulation intensity lacks clear scientific justification (Turi et al., 2022). Additionally, with this uniform intensity setting, TMS successfully modulated statistical learning in a prior study (Ambrus et al., 2020). TMS coil positioning followed the international 10-20 electroencephalography (EEG) system using an EEG cap – this method can be used with a 90% accuracy to position the coil over the targeted area (Herwig et al., 2003). The center of the coil was placed at the location of the F3 electrode for left DLPFC stimulation and at F4 for right DLPFC stimulation (Brodmann 9) throughout the entire stimulation period. In the case of bilateral DLPFC stimulation, the coil was placed at the F3 location for the first half of the stimulation (5 minutes, 300 pulses) and then moved to F4 for the second half (see Figure 3b). The order of stimulation for the hemispheres was counterbalanced across participants in the Bilateral group. For sham stimulation, the coil was tilted 90 degrees away from the skull, thus, the participants could hear the noise made by the machine, but it had no effect on the brain functioning.

**Figure 3.**
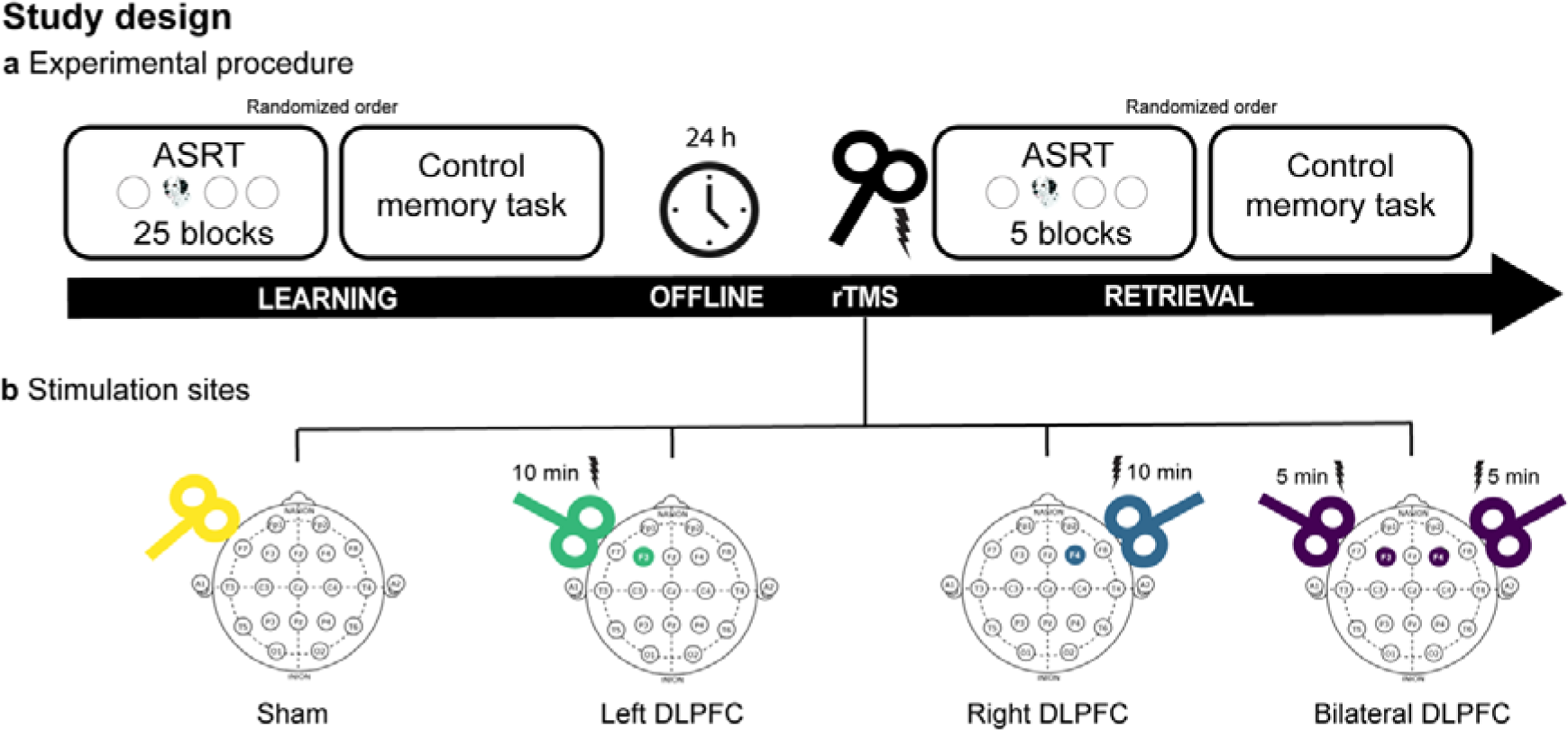
Study design. **a** The study spanned two experimental days. On the first day, participants practiced the ASRT task through 25 learning blocks and performed the learning phase of the PALT, then a 24-hour offline period ensued. On the second day, participants received 1 Hz rTMS for 10 minutes. Immediately after rTMS administration, participants performed 5 blocks of the ASRT task and the recall phase of the PALT. The order of the two tasks was counterbalanced between participants on both days. **b** Stimulation sites for the four groups: coil was tilted by 90 degrees in the sham group, F3 was stimulated for 10 minutes in the Left DLPFC group, F4 was stimulated for 10 minutes in the Right DLPFC group, and F3 and F4 were sequentially stimulated for 5-5 minutes in the Bilateral Group.

### Procedure

The study spanned two experimental days, during which participants engaged in tasks in a well-lit, quiet environment. The initial day involved participants performing the ASRT task across 25 blocks, lasting approximately 25-35 minutes, to acquire the 8-element probabilistic sequence, and the learning phase of the PALT, lasting approximately 10 minutes (Learning session). In the case of the ASRT task, the participants were unaware of the learning nature of the task. Additionally, on the first day, participants completed the Counting Span task to ensure that the four experimental groups did not differ in baseline cognitive functions (see Table 1). After a 24-hour offline period, the second day involved the administration of rTMS and the retest of participants’ statistical and declarative knowledge (Retrieval session). The rTMS procedure, lasting 10 minutes, was immediately followed by the ASRT task comprising 5 blocks using the same alternating sequence practiced on the previous day, or the recall phase of the PALT. The order of the statistical and declarative learning tasks was counterbalanced between and within participants in the two sessions (see Figure 3a).

### Statistical analysis

#### ASRT

Trills (such as 1-2-1) and repetitions (such as 1-1-1) were omitted from the analysis because subjects might exhibit inherent response patterns for these trial types (Soetens et al., 2004). Additionally, trials with RTs below 100 ms and those exceeding three SDs above the mean RT were excluded, as they are unlikely to represent valid reactions. Trials with incorrect reactions (misses) were also removed.

Statistical analysis was performed in R. Linear mixed models (LMMs) were fitted on block-wised mean RT data with the *mixed* function of the *afex* package, separately for the Learning Session and Retrieval Session. Trial Type (high-vs. low-probability), Group (Left, Right, Bilateral, Sham) and Block (Learning Session: 1-25; Retrieval Session: 1-5) were included as fixed factors. Subject factor was included as random intercept, as well as by-participant correlated slopes for the Block factor. To assess the significant factors influencing the model’s quality, we conducted a Likelihood ratio test by utilizing the *anova* function in R. This test compared the likelihoods, as indicated by the Akaike Information Criteria (AIC) in LMMs, between two or more models. The model with the lowest AIC was considered the best model with the highest information gain (Bozdogan, 1987). Estimated marginal means were computed with the *emmeans* R package. Alpha level of .05 were applied to all analyses. If necessary, Bonferroni-correction was performed on post-hoc paired comparisons.

#### PALT

Three learning indices could be distinguished based on the answers of the participants. Item memory index was calculated by subtracting the ratio of incorrect Old-Old responses to New-New pairs (false alarm) from the ratio of responses indicating recognition of Old-Old rearranged pairs (hit rate). Association learning index was quantified by subtracting the ratio of responses indicating recognition of rearranged Old-Old responses (hit rate) from the ratio of responses indicating recognition of original Old-Old responses (hit rate). Finally, the recollection index was defined by subtracting the ratio of incorrect Old-Old original responses to Old-Old rearranged pairs (false alarm) from the ratio of responses indicating recognition of Old-Old original pairs (hit rate). To compare the three PALT learning indices between the four groups, one-way analyses of variances (ANOVAs) were conducted.

## Results

### Comparable statistical learning performance in the four groups in the Learning session

The best model included Trial Type, Block, and Group as fixed factors, where Block was also added as a by-participant random slope factor (see Supplementary Table 1). A main effect of Trial Type was found, with high-probability trials showing faster RTs than low-probability trials, thus statistical learning occurred among all participants (*F*(1, 4840) = 278.76, *p* < .001). The interaction between Trial Type and Block revealed a progressive improvement in statistical learning with gradually increasing difference between high- and low-probability triplets (*F*(1, 4840) = 33.62, *p* < .001). However, the lack of interaction between Group and Trial Type, as well as between Group and Trial Type and Block, ensured that the four groups did not differ in statistical learning (*F*(3, 4840) = 0.45, *p* = .714), or in the course of statistical learning (*F*(3, 4840) = 0.32, *p* = .814) prior to the stimulation (see Figure 4a-b). The main effect of Block revealed decreasing RTs throughout the task, indicating the gradual improvement of visuomotor performance (*F*(1, 97) = 321.79, *p* < .001). The four groups did not differ in visuomotor performance, as Group main effect (*F*(3, 97) = 0.05, *p* = .987) and interaction between Group and Block (*F*(3, 97) = 2.09, *p* = .106) were not found to be significant.

**Figure 4.**
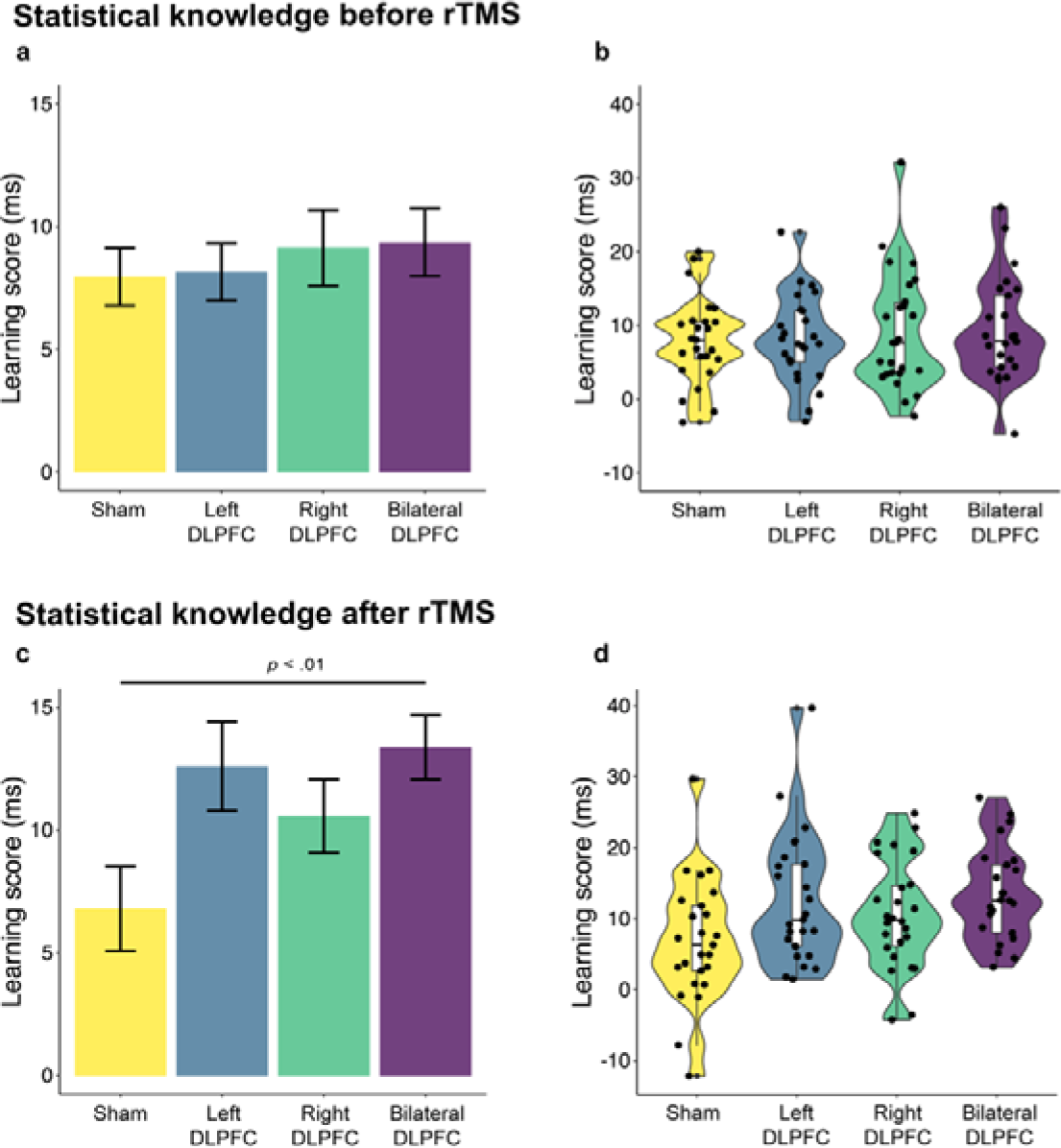
Statistical knowledge before and after rTMS. **a** The y-axis shows the mean statistical learning scores (RT difference between high- and low-probability trials in ms) of the four groups of the learning session. The DLPFC groups did not differ from the Sham group in statistical learning. Error bars represent stand error. **b** Individual statistical learning scores in the four groups. **c** The y-axis shows the mean statistical learning scores (RT difference between high- and low-probability trials in ms) of the four groups of the retrieval session. The Bilateral DLPFC group outperformed the Sham group in the retrieval of statistical knowledge (*p* < .01). Error bars represent standard error. **d** Individual statistical learning scores in the four groups.

### Enhanced retrieval capacity of statistical knowledge after bilateral DLPFC stimulation in the Retrieval session

The best model regarding the Retrieval session included Trial Type and Group as fixed factors, as well as their interaction, and Block as a by-participant random slope factor (see Supplementary Table 2). Intact statistical learning was evidenced by the main effect of Trial Type, with greater speed for high-probability triplets compared to low-probability ones (*F*(1, 804) = 199.10, *p* < .001). The absence of Group main effect indicated that the visuomotor performance of the four groups was comparable (*F*(3, 97) = 0.42, *p* = .737). Nonetheless, the interaction between Group and Trial Type revealed a difference in overall statistical learning between the four groups (*F*(3, 804) = 3.62, *p* = .013). According to post-hoc Welch’s t-tests, two stimulation groups demonstrated better statistical learning performance compared to the Sham group: the Bilateral DLPFC group (*t*(44.835) = 3.04, *p* < .01) and the Left DLPFC group (*t*(47.839) = 2.32, *p* < .05), which remained significant only in the Bilateral DLPFC group after correction for multiple comparisons (see Figure 4c-d).

### Intact recall capacity on the control memory in the Retrieval session

The four groups were found to be comparable with regard to declarative performance. Item memory index was similar in the four groups (*F*(3, 97) = 1.017, η*^2^p* = 0.031, *p* = .389). Association learning index did also not differ in the four groups (*F*(3, 97) = 1.298, η*^2^p* = 0.039, *p* = .279). Finally, recollection index was similar in the four groups (*F*(3, 97) = 0.240, η*^2^p* = 0.007, *p* = .869).

## Discussion

In this study, we investigated a pivotal aspect of predictive processing: the function of the DLPFC in retrieving predictive models. Here, we aimed to fill this gap by administering low-frequency rTMS over the left, right, and bilateral DLPFC before retesting participants’ statistical knowledge acquired 24 hours prior. Our results indicate that disrupting the DLPFC enhances retrieval capacity, particularly if we stimulate both hemispheres. Since general visuomotor performance (speed regardless of trial probability) and the control memory task remained unaffected by rTMS intervention, this boosting effect is presumed to be specific to statistical learning. These findings suggest that less DLPFC involvement not only aids in the acquisition as found by Ambrus et al. (2020) but also the retrieval of temporally distributed predictable patterns, providing further evidence for the PFC’s switching role between competitive memory processes.

Our findings align with previous research demonstrating improved statistical learning following DLPFC suppression by rTMS. Smalle et al. (2017; 2022) applied continuous theta burst stimulation (cTBS), and they showed increased learning of linguistic sequences after cTBS over the left DLPFC. Galea et al. (2010) stimulated the left and right DLPFC in separate groups with cTBS immediately after training on a non-linguistic motor sequence learning task. They found improved learning capacity eight hours later in both groups compared to control, with greater improvement after right DLPFC stimulation than after left DLPFC. Similarly, applying cTBS over the right DLPFC after training led to offline improvement in non-linguistic statistical learning, whereas its facilitatory counterpart did not have a boosting effect (Tunovic et al., 2014). These findings confirm the notion that the PFC plays a crucial role in acquiring statistical regularities and now, supplemented by our findings, also in the retrieval processes.

All those studies that found increased statistical learning capacity on cognitive depletion through DLPFC inhibition interpret their findings in the context of competitive memory processes (Ambrus et al., 2020; Galea et al., 2010; Smalle et al., 2017; Smalle et al., 2022). In the traditional taxonomy based on consciousness, statistical learning falls under non-declarative (or procedural) memory, which encompasses the implicit, habitual acquisition of rigid associations (Henke, 2010; Squire, 1992). The opposing memory system is declarative memory, associated with explicit, goal-directed learning of flexible associations (e.g., episodic memory). The two memory systems rely on distinct neural substrates: the former is supported by basal ganglia and striatum, while the latter is supported by the PFC and medial temporal lobe (MTL) (Squire, 1992; Henke, 2010). In this framework, PFC is hypothesized to promote cognitive control-dependent declarative memory while inhibiting habitual, associative learning processes (Janacsek et al., 2012; Juhasz et al., 2019; Smalle & Möttönen, 2023). In this scenario, less DLPFC involvement translates to more available cognitive resources for statistical learning.

Here, we propose a new hypothesis regarding the mechanism by which PFC mediates between memory processes. In a contemporary framework, opposing memory types are not supported by distinct brain structures but share anatomical backgrounds (A. C. Schapiro et al., 2014, 2017; Sherman, Turk-Browne, et al., 2024). Thus, the hippocampus (MTL), traditionally associated with declarative memory, is now suggested to play a role in both specific episodic memories and more general statistical regularities (Covington et al., 2018; Henin et al., 2021; Schapiro et al., 2014; Schapiro & Turk-Browne, 2015; Sherman, Turk-Browne, et al., 2024). On the other hand, the striatum, traditionally linked to skill acquisition, is implicated in both goal-directed and habitual behavior (Sherman et al., 2024). Within this framework, the PFC is proposed to function as a top-down controller that modulates the activity of both the hippocampus and the striatum to shift between habits and goals (Oehrn et al., 2018; Sherman et al., 2024; Williams, 2023). This hypothesis gains further support from the results of an intracranial EEG study, where directed forgetting was associated with decreased hippocampal activation determined by increased DLPFC activation (Oehrn et al., 2018). Moreover, a neuroimaging study demonstrated that statistical learning performance could be predicted by the integrity of the tract between the PFC and hippocampus (Bennett et al., 2011). In light of these insights, we propose that the PFC acts as a flexible controller, finely tuning the expression of different memory subsystems to suit task demands. Nevertheless, future studies utilizing neuroimaging methods should elucidate the pathway by which reduced PFC involvement contributes to enhanced statistical learning and retrieval.

However, apparent contradictions arise regarding lateralization. In our research, the retrieval performance of the left and right DLPFC groups was similar to the retrieval capacity of the bilateral DLPFC group, yet it did not surpass that of the sham group, even though other studies found unilateral stimulation to be effective (Smalle et al., 2017; Tunovic et al., 2014). One possible reason for this discrepancy could be the different stimulation protocols, as rTMS and TBS could exert different effects on statistical learning (Glinski, 2021; Verwey et al., 2022). Furthermore, the complexity of statistical information present in the input could be a significant factor. In our study, right DLPFC stimulation was the least effective among the three active stimulation conditions in retrieval capacity modulation, whereas other studies found right hemisphere stimulation to be effective (Tunovic et al., 2014), even more effective than stimulation of the left DLPFC (Galea et al., 2010). However, those studies used deterministic sequences, which, unlike the probabilistic sequences we utilized, have a much simpler structure and are less effective in accurately reflecting real-life statistical learning processes. In our study, the left DLPFC group, similar to the bilateral DLPFC group, achieved significantly better retrieval compared to the sham group, but this performance difference disappeared after correction for multiple comparisons. In another study, where statistical learning of linguistic sequences was investigated, inhibitory stimulation of the left DLPFC also led to better learning. These findings suggest that more complex sequences with higher ecological validity are likely to rely more on the left hemisphere. Further studies are needed to support the lateralization of statistical learning and retrieval in the PFC.

Our finding of the superiority of bilateral stimulation over unilateral stimulation is highly consistent with that of a previous study (Ambrus et al., 2020). They administered 1 Hz rTMS to the DLPFC bilaterally between the learning blocks and observed subsequently enhanced learning capacity of probabilistic regularities compared to the sham group. The authors suggested that sequential bilateral stimulation, where the same rTMS protocol is successively applied to both hemispheres, may prevent potential interhemispheric compensatory mechanisms. In our study, the statistical knowledge following left and right DLPFC stimulation approached that of bilateral stimulation, albeit not to the extent that it surpassed the performance of the Sham group. This is likely due to the unstimulated hemisphere compensating for the other’s function thereby neutralizing the stimulation effect. Another important consideration is that studies where only one hemisphere was stimulated typically involve one-handed versions of motor learning tasks (Szücs-Bencze et al., 2023). For studies using two-handed tasks, the bilateral approach is generally more advisable.

The DLPFC encompasses a large area in the brain with potentially differing functions across subregions, making precise localization crucial when investigating its role in cognitive functions. In order to paint a more accurate picture of the role of the PFC and confirm the pathway through which reduced PFC activation exerts its beneficial effect on statistical learning and retrieval, it would be of paramount importance to employ electrophysiological or neuroimaging techniques (e.g., functional magnetic resonance imaging) combined with NIBS. Future studies should aim to replicate this research using methodologies like psychophysiological interaction analysis (PPI) and involving functional connectivity toolboxes like CONN (Whitfield-Gabrieli & Nieto-Castanon, 2012), which would allow the confirmation of whether the functional link between distant brain areas (e.g., PFC and hippocampus) is responsible for statistical learning performance (O’Reilly et al., 2012).

In conclusion, we demonstrated the functional role of DLPFC in retrieving previously acquired statistical knowledge a key component in predictive coding and processing. The most significant enhancement in retrieval capacity occurred with bilateral stimulation, suggesting that suppressing the DLPFC is particularly beneficial when interhemispheric compensatory mechanisms are limited. These findings enrich our understanding of the PFC’s role in predictive processing and switching between competing memory systems and have implications for cognitive enhancement strategies.

## Acknowledgments

This research was supported by the National Brain Research Program (Project NAP2022-I-2/22) and the Chaire de Professeur Junior Program by INSERM and the French National Grant Agency (ANR-22-CPJ1–0042–01ANR) (to N. D.). Prepared with the support of the Gedeon Richter’s Talentum Foundation established by Gedeon Richter Plc. (to Sz-B. L.).

