## Supplementary Table 1, 2 for "Enhancing Retrieval Capacity of the Predictive Brain through Dorsolateral Prefrontal Cortex Intervention"

**Supplementary Table 1.** Linear mixed model analysis of mean RTs of Learning session

| Fixed effects | *b* | *SE b* | *95% CI* | *t* | *p* |
| --- | --- | --- | --- | --- | --- |
| (Intercept) | 359.62 | 3.97 | [351.83, 367.41] | 90.52 | < .001 |
| Bilateral DLPFC | 0.67 | 6.90 | [-12.86, 14.20] | 0.09 | .92 |
| Left DLPFC | 1.57 | 6.90 | [-11.95, 15.11] | 0.22 | .81 |
| Right DLPFC | 0.08 | 6.90 | [-13.44, -13.61] | 0.01 | .99 |
| Block | -13.26 | 0.73 | [-14.71, -11.81] | -17.93 | < .001 |
| Trial Type | -4.32 | 0.25 | [-4.83, -3.81] | -16.69 | < .001 |
| Bilateral DLPFC × Block | 1.91 | 1.28 | [-0.60, 4.43] | 1.49 | .13 |
| Left DLPFC × Block | -2.67 | 1.28 | [-5.19, -0.15] | -2.08 | .03 |
| Right DLPFC × Block | -0.77 | 1.28 | [-3.29, 1.74] | -0.60 | .54 |
| Bilateral DLPFC × Trial Type | 0.34 | 0.45 | [-0.53, 1.23] | 0.77 | .44 |
| Left DLPFC × Trial Type | -0.35 | 0.45 | [-1.23, 0.53] | -0.78 | .43 |
| Right DLPFC × Trial Type | 0.24 | 0.45 | [-0.63, 1.13] | 0.55 | .58 |
| Block × Trial Type | -1.50 | 0.25 | [-2.01, -0.99] | -5.79 | < .001 |
| Bilateral DLPFC × Block × Trial Type | 0.05 | 0.45 | [-0.82, 0.93] | 0.12 | .90 |
| Left DLPFC × Block × Trial Type | 0.34 | 0.45 | [-0.53, 1.22] | 0.76 | .44 |
| Right DLPFC × Block × Trial Type | -0.04 | 0.45 | [-0.92, 0.83] | -0.10 | .91 |
| Random effects |  |  |  |  |  |
| σ^2^ | 338.86 |  |  |  |  |
| τ00_Subject_ | 1586.93 |  |  |  |  |
| τ11_Subject.Block_ | 48.42 |  |  |  |  |
| ρ01_Subject_ | -0.38 |  |  |  |  |
| ICC | 0.83 |  |  |  |  |
| N_Subject_ | 101 |  |  |  |  |
| Observations | 5050 |  |  |  |  |
| Marginal R^2^/Conditional R^2^ | 0.93/0.84 |  |  |  |  |

| Fixed effects | *b* | *SE b* | *95% CI* | *t* | *p* |
| --- | --- | --- | --- | --- | --- |
| (Intercept) | 319.02 | 3.06 | [313.00, 325.04] | 104.03 | < .001 |
| Bilateral DLPFC | 5.34 | 5.32 | [-5.11, 15.80] | 1.00 | .31 |
| Left DLPFC | -3.51 | 5.32 | [-13.97, 6.93] | -0.66 | .50 |
| Right DLPFC | 0.71 | 5.32 | [-9.74, 11.17] | -0.13 | .89 |
| Trial Type | -5.41 | 0.38 | [-6.16, -4.66] | -14.11 | < .001 |
| Bilateral DLPFC × Trial Type | 2.01 | 0.66 | [0.70, 3.32] | 3.02 | .003 |
| Left DLPFC × Trial Type | -1.26 | 0.66 | [-2.57, 0.04] | -1.89 | .058 |
| Right DLPFC × Trial Type | -0.88 | 0.66 | [-2.19, 0.42] | -1.32 | .18 |
| Random effects |  |  |  |  |  |
| σ^2^ | 148.73 |  |  |  |  |
| τ00_Subject_ | 967.99 |  |  |  |  |
| τ11_Subject.Block_ | 19.88 |  |  |  |  |
| ρ01_Subject_ | -0.25 |  |  |  |  |
| ICC | 0.87 |  |  |  |  |
| N_Subject_ | 101 |  |  |  |  |
| Observations | 1010 |  |  |  |  |
| Marginal R^2^/Conditional R^2^ | 0.03/0.87 |  |  |  |  |

**Supplementary Table 2.** Linear mixed model analysis of mean RTs of Retrieval session
